# Effects of acute lying and sleep deprivation on the behavior of lactating dairy cows

**DOI:** 10.1101/549378

**Authors:** Jessie A. Kull, Katy L. Proudfoot, Gina M. Pighetti, Jeffery M. Bewley, Bruce F. O’Hara, Kevin D. Donohue, Peter D. Krawczel

## Abstract

The objective was to determine the effects of sleep or lying deprivation on the behavior of dairy cows. Data were collected from 8 multi- and 4 primiparous cows (DIM = 199 ± 44 (mean ± SD); days pregnant = 77 ± 30). Using a crossover design, each cow experienced: 1) sleep deprivation implemented by noise or physical contact when their posture suggested sleep, and 2) lying deprivation imposed by a grid placed on the pen floor. One day before treatment (baseline), and treatment day (treatment) were followed by a 12-d washout period. Study days were organized from 2100 to 2059. During habituation (d −3 and −2 before treatment), baseline (d −1), and trt (d 0), housing was individual boxstalls (mattress with no bedding). After treatment, cows returned to sand-bedded freestalls for a 7-d recovery period (d 1 to 7) where data on lying behaviors were collected. Daily lying time, number lying bouts, bout duration, and number of steps were recorded by dataloggers attached to the hind leg of cows throughout the study period. Data were analyzed using a mixed model in SAS including fixed effects of treatment (sleep deprivation vs. sleep and lying deprivation), day, and their interaction with significant main effects separated using a PDIFF statement (*P* ≤ 0.05). Interactions between treatment and day were detected for daily lying time and the number of bouts. Lying time was lower for both treatments during the treatment period compared to baseline. Lying time increased during the recovery period for both lying and sleep deprived cows. However, it took 4 d for the lying deprived cows to fully recover their lying time after treatment, whereas it took the sleep deprived cows 2 d for their lying time to return to baseline levels. Results suggest that both sleep and lying deprivation can have impact cow behavior. Management factors that limit freestall access likely reduce lying time and sleep, causing negative welfare implications for dairy cows.

## INTRODUCTION

Lying time is critical for biological function for dairy cows, however, there are various factors on-farm that reduce a cow’s ability to lie down or influence how she uses her lying space. Management factors such as overstocking (Krawczel et al., 2012) or heat stress (Cook et al., 2007) decrease lying time, either by reducing access to lying spaces or altering the cow’s motivation to lie down. Additionally, facility factors such as bedding type (Fregonesi et al., 2007a) and stall design (Fregonesi et al., 2009), can influence how a cow uses a lying stall. The impact of facility design on lying time has been well-studied, but less is known about the quality of rest cows are able to maintain while lying down. A measurement of rest quality in cattle and other species is sleep, but very little research has assessed sleep in dairy cattle.

Within their time budget, dairy cows lie down between 11 and 13 h/d in confinement housing systems (Tucker and Weary, 2004, Jensen et al., 2005, Ito et al., 2009). A portion of lying time is spent sleeping. Sleep is divided into two main vigilant states: non-rapid eye movement (NREM) sleep, and rapid eye movement (REM) sleep (Irwin, 2015). However, drowsing in some animals is observed, and described as an intermediate state between wake and NREM sleep (Ruckebusch, 1972). Dairy cows sleep for about 4 h/d, in short 3 to 5 minute bouts throughout the day (Ternman et al., 2012). Specifically, they spend 3 h/d in NREM sleep, 30 to 45 min/d in REM sleep, and 8 h/d drowsing (Ruckebusch, 1972). Cows can also drowse and engage in some NREM sleep when forced to stand, though, this is not normally observed (Ruckebusch, 1972). All sleep states cannot be achieved while standing; a recumbent position must be assumed for cows to engage in REM sleep (Ruckebusch, 1972). Therefore, any loss of lying time has the potential to alter the time cows spend sleeping.

Research has found that lying time is an important behavior for dairy cows. When given the choice, cows relinquish other activities such as feeding and socializing to spend more time lying down (Munksgaard et al., 2005). During a 2 or 4 h lying deprivation period, cows stomped their feet, shifted their weight, and head butted neighboring cows (Cooper et al., 2007). These behaviors were consistently observed during lying deprivation periods of 22 h/d for two weeks (Ruckebusch, 1974), and 3 h/d for one week (Metz, 1985). Collectively, this suggests cows are likely expressing frustration, restlessness and lack of comfort during this time. While lying time was reduced in these studies, some degree of sleep deprivation is likely imposed as well, because cows cannot engage in REM sleep while standing (Ruckebusch, 1974). Therefore, it is not known if the effects observed during lying deprivation are solely a result of lying deprivation, or the cumulative effects of lying and sleep deprivation.

A reduction in lying time has been found to affect cow productivity. For example, Bach et al. (2008) found that as free stall access decreased, milk production was reduced, suggesting lying time plays a critical role in milk yield. With each additional hour of lying time, the cow produces 0.91 to 1.59 kg of milk per day (Grant, 2004). The impact of sleep loss on milk yield in dairy cattle is unknown, however there is research in other species to suggest that sleep may be important for lactation. For example, growth hormones and prolactin, which are key hormones associated with milk production, are decreased during sleep deprivation in humans (Davidson et al., 1991). To date, research has evaluated the effects of lying deprivation, but did not account for sleep deprivation. Thus, the primary objective of this study was to determine the effect of a 24-h period of sleep and/or lying deprivation on the lying behavior of dairy cows during deprivation and during a 7 d recovery period. A second objective was to determine the impact of a 24-h lying deprivation on sleep states in cattle. Thirdly, we determined the impact of sleep and/or lying deprivation on the milk production of dairy cows.

## MATERIALS AND METHODS

### Animals, Housing and Management

This study was conducted at the University of Tennessee’s Little River Animal and Environmental Unit (Walland, TN) during April and May 2016. Mid to late-lactation Holstein dairy cows (n = 12) were enrolled based on DIM (DIM = 199 ± 44) and days pregnant (77 ± 30 d). Cows were milked twice daily starting at 0700 and 1730 h in a double-8 herringbone milking parlor (BouMatic, Madison, WI). Cows are normally housed in deep-bedded sand freestall pens. During the study period, cows were housed individually in a 4.11 × 3.32 m pen with a mattress and no bedding. Visual and olfactory contact between cows was possible for enrolled cows throughout the duration of the treatment phase. Individually housing in this manner facilitated the use of electrophysiological equipment to assess vigilant state and lying deprivation. Pens were thoroughly scrubbed with chlorhexidine solution (Durvet Inc., Blue Springs, MO) every morning at 0700 h when cows were being milked. Fecal matter was removed manually throughout the day to maintain pen and cow hygiene. Fresh water and a TMR were available ad libitum. The TMR was comprised of 60% corn silage, 25% pelleted premix grain concentrate, 12% small grain silage, and 3% dry hay. All procedures described were approved by the University of Tennessee Institutional Animal Care and Use Committee.

### Enrollment Criteria

From the cows meeting the selection criteria for DIM and pregnancy, a final group of 12 cows were selected using white blood cell count (WBC ≤ 12.6), and temperament. Blood samples from the target population of cows, were taken via the coccygeal vein and WBCs were analyzed to ensure cows were below the accepted threshold of 12.6 cell/mL as described by Schalm (1961), indicating the cows were not experiencing any prior illness. Thus, cows enrolled in the study were considered healthy. Temperament was evaluated using an approachability and brush test. For the approachability test, a researcher slowly approached the cow with one arm extended, and observed the cow’s reaction (Lensink et al., 2003). Cows were scored based on the 1 to 4 scale described by Lensink et al. (2003), with 1 being defined as the cow allowing physical contact, and 4, the cow strongly withdrew from the researcher (Table 1). If the cow remained still and allowed physical contact or approached the researcher (a score of 1 or 2), the cow was considered suitable for the study. The brush test used was slightly modified from the brush test described by Ternman et al. (2014), where cows were restrained in pen headlocks, instead of free roaming. Cows were scored based on a 1 to 4 scale, similar to the scale defined by Lensink et al. (2003) (Table 1). For this test, the cow’s head and neck area were brushed, particularly where the EEG equipment would be placed (Lensink et al., 2003, Ternman et al., 2014). If the cow did not pull away, or only slightly withdrew when brushing occurred (a score of 1 or 2), she was considered an acceptable candidate for the study. In total, 14 cows met the criteria for the temperament tests, however, 2 cows were removed because their WBC count exceeded the accepted 12.6 cell/mL threshold.

**Table 1.**
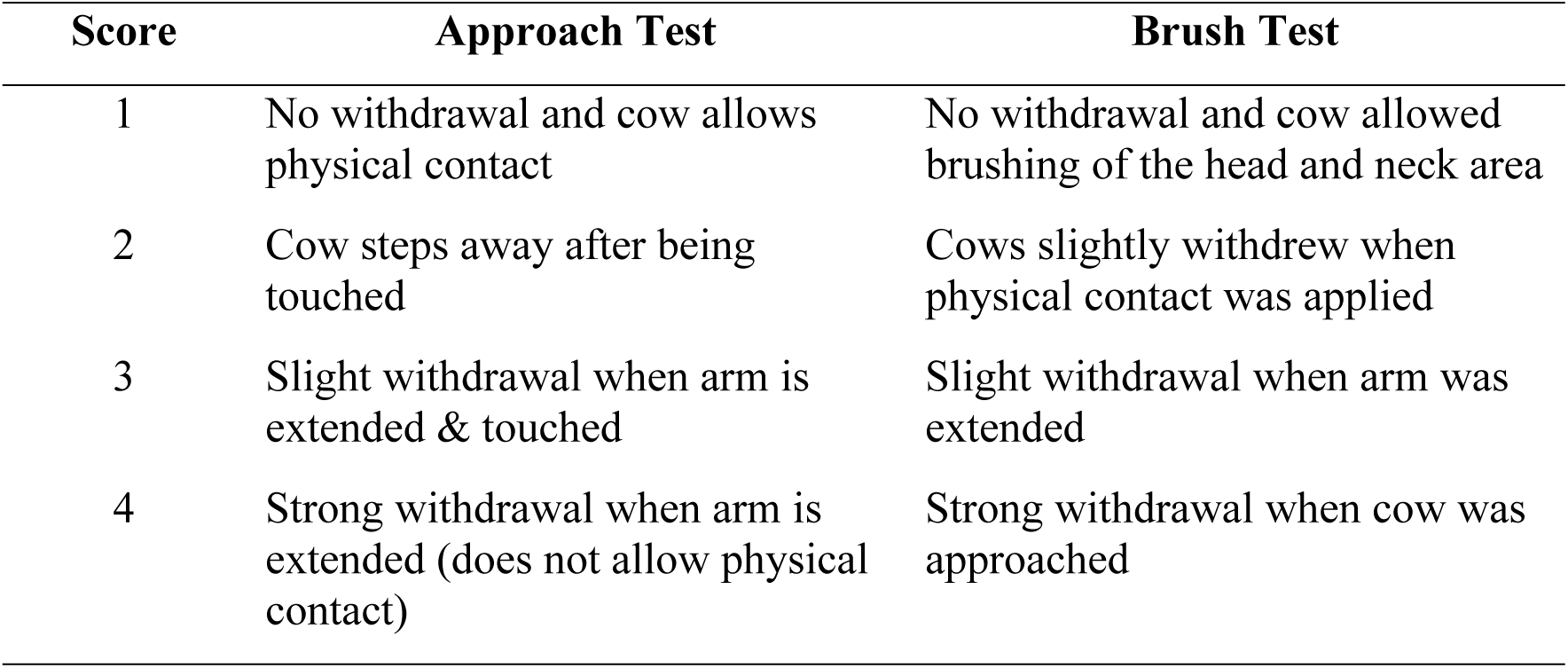
Approachability and brush test scoring guide using a 1 through 4 scale modified from Lensink et al. (2003). Cows were deemed acceptable for the study if they were scored a 1 or 2.

### Treatments

Treatments were implemented using a crossover design with rolling enrollment. The study design progressed from a habituation (−3 d, −2 d), baseline (−1 d), treatment (0 d) and recovery (1 – 7 d) period, with a 12-d washout period between treatments. Each ‘day’ represented a 24 h period from 2100 to 2059 to facilitate the treatments used in the study. Cows were housed in the individual pens during the habituation, baseline and treatment phases, and were moved back to the group free-stall pen for the recovery period. Because cows were moved to an unfamiliar pen at the start of the study, a 2-d habituation period was provided to allow cows to adapt to their new environment. When cow were regrouped into a novel pen (von Keyserlingk et al., 2008), or trained to use a robotic milking system (Jacobs and Siegford, 2012), it only took the cows 2 d to habituate, suggesting our 2-d habituation period was sufficient. Additionally, the mattress bedded pens the cows were placed in were only 8 m away from their home pen, so visual, and olfactory contact were maintained.

All cows experienced two treatments: 1) a 24-h lying deprivation period, and 2) a 24-h sleep deprivation. After the treatment phase, cows returned to their home deep-bedded sand freestall pen for a 7-d recovery period and a 12-d washout period before returning to an individual pen for their second treatment (whichever treatment they did not experience first). The order of treatments was randomized for each cow.

#### Lying Deprivation

The 24-h lying deprivation period was implemented using a wooden girl placed on the pen floor, preventing cows from assuming a recumbent position. The wooden grid was based on a design by Schütz et al. (2008) which prevented cows from lying during times of heat stress.

#### Sleep Deprivation

During the 24-h sleep deprivation period, cows were allowed to lie down, but were continuously monitored to ensure cows remained awake and alert. If a cow’s posture suggested the onset of sleep, such as her eyes closed and neck relaxed, the cow would be touched to keep her awake as described by Ledoux et al. (1996) who used this method in cats. Gentle handling or touching was used because it implemented deprivation, but likely did not induce a stress response that would be caused by the method of deprivation (Graves et al., 2003).

### Behavioral Data

#### Lying Behaviors

IceTag dataloggers (IceRobotics Ltd., Edinburgh, Scotland) were attached to the hind leg of each cow during milking two days before to the start of the study to allow for habituation (MacKay et al., 2012). A total of 18 d worth of data were collected and analyzed from the IceTags for each cow (1 baseline day, 1 treatment day and 7 recovery days for each of the two treatments). The IceTags collected daily lying times (h/d), lying bout frequency (number/d), lying bout length (min/bout), and total steps (number/d) (McGowan et al., 2007).

#### Electrophysiological data

During the baseline period, cows were fitted with the electrophysiological equipment that collected electroencephalographic (EEG), electrooculography (EOG), and electromyography (EMG) data (EEG; BioRadio, Great Lakes Neurotechnologies, Cleveland, OH). Cows were restrained in the headlock of the experimental pen for placement of the electrophysiological equipment. Hair at the location of each electrode was shaved using 40 blade clippers (Andis, Sturtevant, WI) and wiped clean with alcohol to ensure sufficient contact. Non-invasive electrodes were plugged into the EEG device and then placed on the cow using Durapore Surgical Tape (3m Healthcare, St. Paul, MN) and adhesive glue (Gorilla Glue Inc., Cincinnati, OH) to secure the electrodes in place. Ten20 EEG conductive paste (Weaver and Company, Aurora, CO) was placed on both sides of the electrodes to help conduct the signal. In total, there were ten electrodes on the cow. Electrode configuration was placed on the head and neck area, and can be further illustrated in Figure 1 (Ternman et al., 2012). During the entire 48 h EEG recordings from the baseline and treatment periods, a researcher was present to monitor the cow and ensure the EEG device, and the electrodes remained in place.

**Figure 1.**
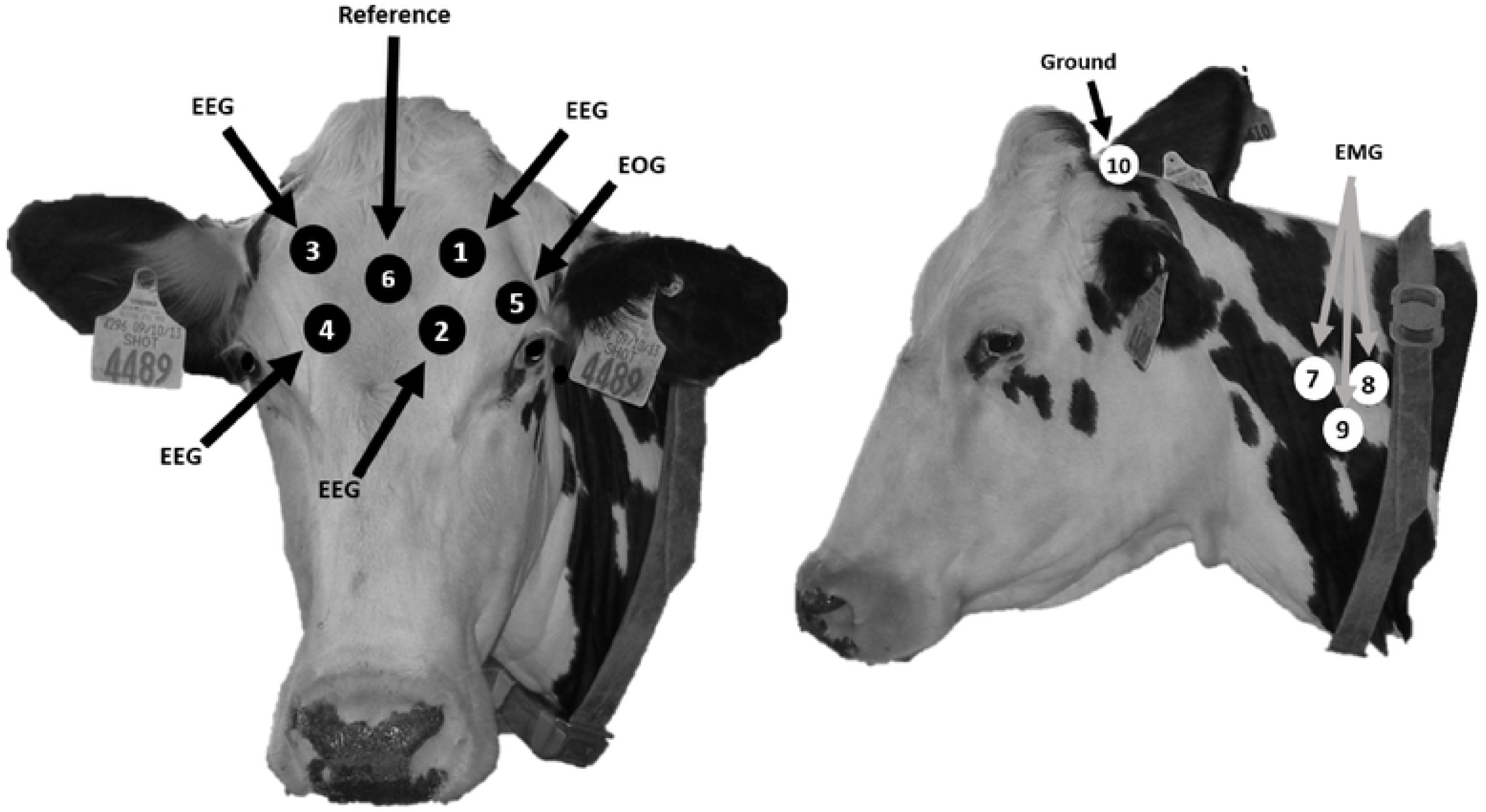
Placement of electrodes as outlined by Ternman et al. (2012)

EEG data were used to determine the amount of REM, NREM and drowsing cows experienced during the baseline and treatment days for a total of 4 d per cow (2 days per cow per treatment).

### Milk Production Data

Cows were milked twice daily starting at approximately 0730 and 1700 h. Milk weights were recorded at each milking on d −2, 2, 3, 4 and 5 relative to treatment (d 0) automatically. The collars that register cows in the parlor were removed during the baseline and treatment days because they interfered with the EEG device. Therefore, milk weights were not recorded during this time. Data from the day before baseline (d −2) when the cow was in the individual pen for habituation was used to represent the baseline period. Milk weights were combined from morning and evening milking to obtain total daily production.

A composite milk sample was collected into a 15mL collection vial during milking on baseline, treatment, and d 2, to monitor fat, protein and somatic cell count (SCC). Morning and evening milk composite data were combined daily for all study days. Milk composite samples were taken automatically via an inline sampler without additional handling of the cow. Samples were stored at room temperature for no more than 48 h before analysis. Milk fat, protein, and somatic cell counts (SCC) were analyzed by the Tennessee Dairy Herd Improvement Laboratory (Knoxville, TN).

### Statistical Analysis

All data were analyzed in SAS 9.4 (SAS Institute, Inc., Carry, NC) using the cow as the experimental unit. Before analysis, all data were screened for outliers and normality. For data that was not normally distributed, such as milk contents, a log transformation was used to normalize all data, and data were reported as back transformed means.

Data were analyzed using the mixed model ANOVA (PROC MIXED) repeated measures. To determine the effect of day and treatment on lying behaviors, models included day (baseline, treatment, and the 7 d recovery period) and treatment (sleep or lying deprivation) as fixed effects, and a day by treatment interaction. To determine if lying behaviors would differ between treatment and baseline periods for cows, specific pair-wise comparisons were made using the PDIFF statement. To determine if lying behavior during the recovery period differed from ‘normal’, we considered d 7 of the recovery period as our reference for ‘normal’. Specific pair-wise were then made between d 7 and every other day in the recovery period for each cow and treatment using the PDIFF statement.

When analyzing the EEG data, a paired t-test was used for all d 1 and 2 comparisons. For the treatment comparisons, a t-test was run for 2 groups assuming equal variance.

To determine the impact of day and treatment on milk production and composition, models included day (d −2, 1, 2, 3, 4, 5 for milk yield and baseline, treatment and d 2 for milk components), treatment (sleep or lying deprivation), and time of sampling (AM or PM sampling) as fixed effects, and a day by treatment interaction. To determine if milk yield differed between baseline (d −2) and the 4 days after treatment, specific pair-wise comparisons were made using the PDIFF statement. Pair-wise comparisons were also made to determine if milk components differed between baseline, treatment and d 2.

To assess the impact of treatment and day on milk composition, Significance was declared a *P* < 0.05 and a trend declared at *P* < 0.10.

## RESULTS

### Lying behavior

When lying behaviors were specifically compared between baseline and treatment days, all behaviors differed during lying deprivation (*P* < 0.05; Table 2). However, there was only a tendency for lying bouts and bout duration to differ during sleep deprivation (*P* < 0.1).

**Table 2.** LS Means and standard errors for lying time, number of lying bouts, lying bout duration, and total steps taken for cows during baseline and treatment days (24 h of lying or sleep deprivation). Cows were housed on mattresses in individual box stalls for both days.

| Variable | Baseline | Treatment | SE | *P*-value |
| --- | --- | --- | --- | --- |
| **Lying Deprivation** |  |  |  |  |
| Lying time, h/d | 8.8 | 1.9 | 0.8 | <.0001 |
| Lying bouts, no/d | 9.6 | 4.1 | 0.8 | <.0001 |
| Lying bout duration, min/bout | 58.9 | 15.3 | 7.3 | <.0001 |
| Steps, no/d | 2422 | 3318 | 261 | <.0001 |
| **Sleep Deprivation** |  |  |  |  |
| Lying time, h/d | 8.6 | 8.4 | 0.7 | 0.71 |
| Number of lying bouts, no/d | 9.0 | 7.6 | 0.8 | 0.07 |
| Lying bout length, min/bout | 61.8 | 72.9 | 7.0 | 0.09 |
| Steps, no/d | 2623 | 2537 | 261 | 0.58 |

When lying behaviors were specifically compared between each day of the recovery period and d 7 (reference), lying time was higher on d 1 compared to d 7 during both lying and sleep deprivation (*P* ≤ 0.006; Figure 2). It took cows 2 and 4 d to recover their lying time after sleep deprivation (P = 0.24) and lying deprivation (*P* = 0.62), respectively.

**Figure 2.**
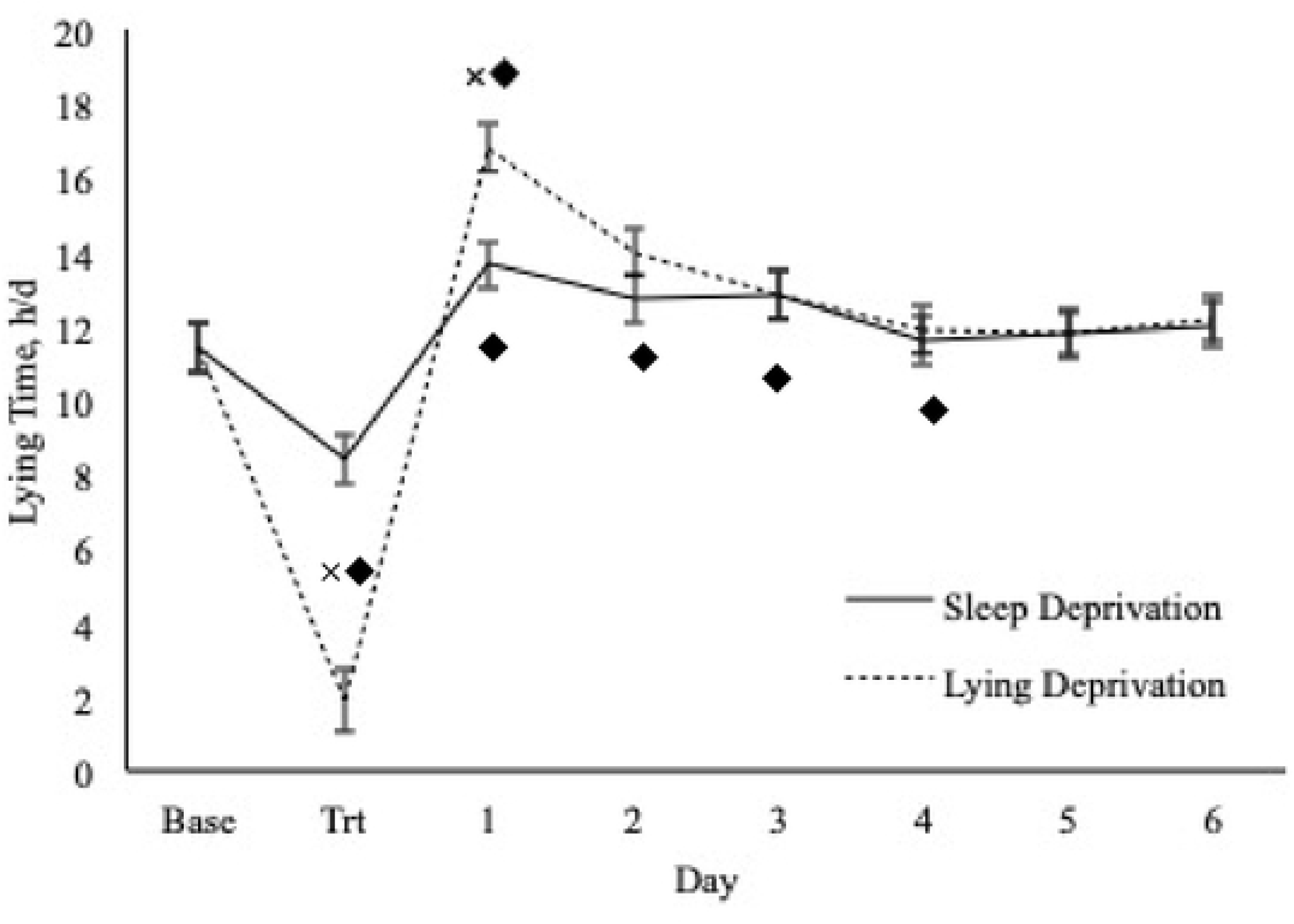
D 7 is used as the baseline period (Base) and d 1 through 6 illustrate the recovery period when cows were returned to their home sand bedded freestall pen. Lying time increased on d 1 for both treatments (trt) (*P* ≤ 0.0003). Lying time did not return to baseline levels until d 5 for the lying deprived cows, and d 2 for the sleep deprived cows (*P* ≥ 0.05). ^**×**◆^Values with a different superscript differ (*P* < 0.05). ^**×**^Indicates across treatment differences, and ^◆^indicates within treatment differences relative to baseline.

Lying bouts did not differ for either treatment on d 1 through 6 compared to d 7 (*P* > 0.05; Table 3). However, there was a tendency for cows to have more lying bouts on d 2 compared to d 7 during lying deprivation (*P* = 0.07). Bout duration was higher on d 1 compared to d 7 for during lying deprivation (*P* < 0.0001). There was a tendency for bout duration to be longer on d 1 compared to d 7 during sleep deprivation (*P* = 0.08; Table 3).

**Table 3.** LS Mean and standard errors of lying bouts, bout duration, and steps for cows that were lying or sleep deprived for 24 h on d 0 during the recovery period (d 1 to 7). All comparisons are made relative to d 7 (reference; the last day of the recovery period). Means with a superscript differed from d 7 (^**^*P* < 0.05 and, ^*^*P* < 0.10).

| Day | Lying Deprivation |  |  | Sleep Deprivation |  |  |
| --- | --- | --- | --- | --- | --- | --- |
|  | Lying bouts (no./d) | Bout duration (min/bout) | Steps (no./d) | Lying bouts (no./d) | Bout duration (min/bout) | Steps (no./d) |
| 1 | 9.7 ± 0.7 | 110.0 ± 6.6** | 1,619 ± 260 | 9.8 ± 0.7 | 89.9 ± 7.0* | 2,010 ± 260 |
| 2 | 10.9 ± 0.7* | 82.1 ± 6.6 | 1,618 ± 260 | 9.3 ± 0.7 | 85.0 ± 7.0 | 1,828 ± 260 |
| 3 | 10.3 ± 0.7 | 80.6 ± 6.6 | 1,686 ± 260 | 10.0 ± 0.7 | 79.3 ± 7.0 | 1,757 ± 260 |
| 4 | 9.8 ± 0.7 | 76.5 ± 6.6 | 2,013 ± 260 | 9.7 ± 0.7 | 72.8 ± 7.0 | 1,925 ± 260 |
| 5 | 10.5 ± 0.7 | 73.1 ± 6.6 | 1,789 ± 260 | 9.9 ± 0.7 | 72.2 ± 7.0 | 1,819 ± 260 |
| 6 | 10.4 ± 0.7 | 74.0 ± 6.6 | 1,805 ± 260 | 9.8 ± 0.7 | 76.3 ± 7.0 | 1,784 ± 260 |
| 7 (ref) | 9.5 ± 0.7 | 76.4 ± 6.6 | 1,729 ± 260 | 10.0 ± 0.7 | 68.7 ± 7.0 | 1,808 ± 260 |

### EEG Data

There was an effect of treatment where NREM sleep decreased from baseline to treatment day (*P* = 0.01). However, there was only a tendency for time spent awake to increase (*P* = 0.09) and REM sleep to decrease (*P* = 0.10). When we combined the lighter sleep states, drowsing, and wake, time spent awake increased from baseline to treatment (*P* = 0.01).

However, when we combined the two deeper sleep states, NREM and REM, there was a decrease between baseline and treatment (*P* = 0.01). When only evaluating lying deprivation, time spent awake increased (*P* = 0.04; Table 4), and time spent in NREM sleep decreased (*P* = 0.02; Table 4). However, there was only a tendency for drowsing (*P* = 0.07; Table 4) and REM sleep (*P* = 0.08; Table 4) to decrease from baseline to treatment. There was no effect of sleep deprivation on any vigilant state (*P* ≥ 0.05; Table 4).

**Table 4.**
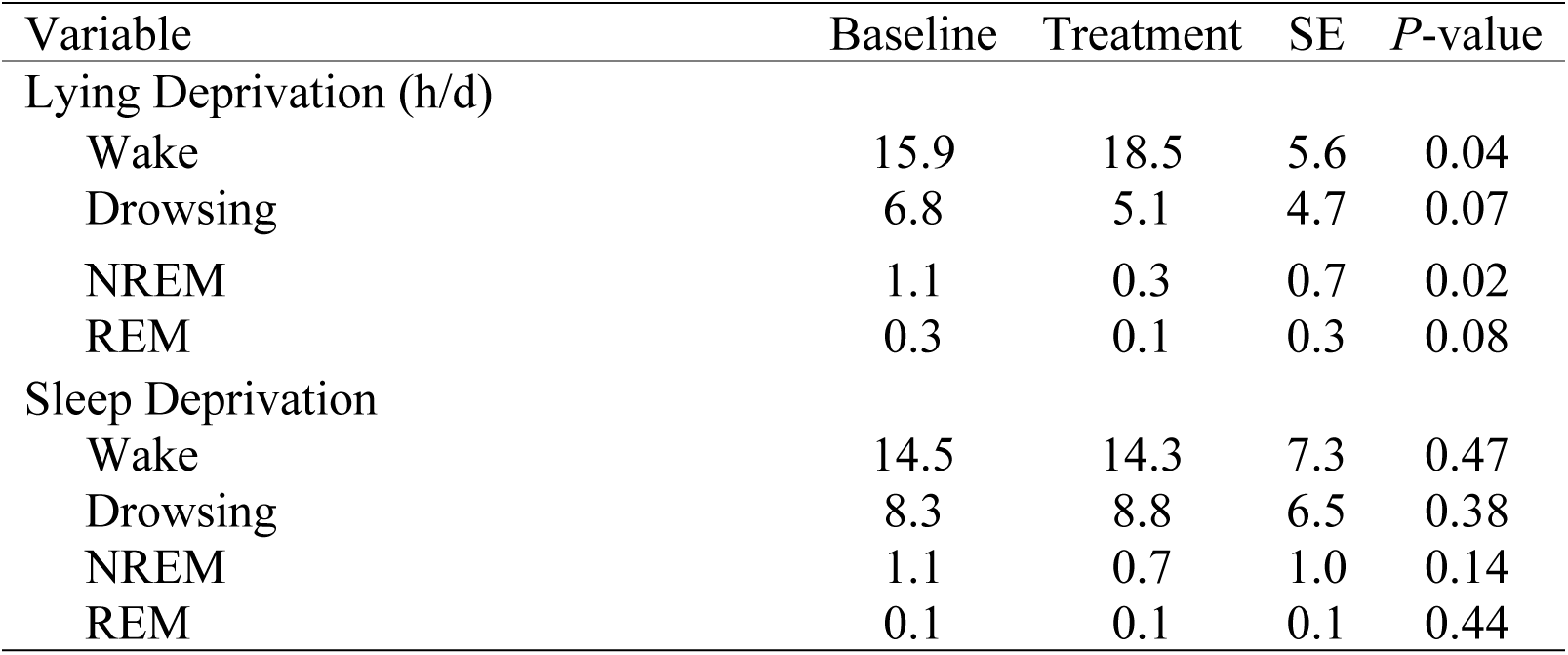
LS Means and standard errors for total hours spent in each vigilant state during baseline and treatment days (24 h of lying or sleep deprivation). Cows were housed on mattress bedding in individual box stalls for both days.

| Variable | Baseline | Treatment | SE | *P*-value |
| --- | --- | --- | --- | --- |
| Lying Deprivation (h/d) |  |  |  |  |
| Wake | 15.9 | 18.5 | 5.6 | 0.04 |
| Drowsing | 6.8 | 5.1 | 4.7 | 0.07 |
| NREM | 1.1 | 0.3 | 0.7 | 0.02 |
| REM | 0.3 | 0.1 | 0.3 | 0.08 |
| Sleep Deprivation |  |  |  |  |
| Wake | 14.5 | 14.3 | 7.3 | 0.47 |
| Drowsing | 8.3 | 8.8 | 6.5 | 0.38 |
| NREM | 1.1 | 0.7 | 1.0 | 0.14 |
| REM | 0.1 | 0.1 | 0.1 | 0.44 |

There was no effect of treatment on the length of sleep bouts (*P* ≥ 0.05). However, there was a tendency for NREM sleep bout length to decrease from baseline to treatment (*P* = 0.07). When evaluating the lying deprivation treatment only, bout length tended to decrease for drowsing and REM sleep (*P* = 0.08; Table 5). However, there was no effect of sleep deprivation on bout length (*P* ≥ 0.05; Table 5).

**Table 5.** LS Means and standard errors for each vigilant state bout length (min/h) during baseline and treatment days (24 h of lying or sleep deprivation). Cows were housed on mattress bedding in individual box stalls for both days.

| Variable | Baseline | Treatment | SE | *P*-value |
| --- | --- | --- | --- | --- |
| Lying Deprivation Bout Length (min/h) |  |  |  |  |
| Wake | 7.2 | 13.6 | 12.0 | 0.04 |
| Drowsing | 2.3 | 1.8 | 0.9 | 0.07 |
| NREM | 1.1 | 0.7 | 0.9 | 0.02 |
| REM | 0.4 | 0.2 | 0.4 | 0.08 |
| Sleep Deprivation Bout Length (min/h) |  |  |  |  |
| Wake | 6.4 | 7.7 | 4.2 | 0.47 |
| Drowsing | 2.7 | 2.8 | 0.6 | 0.38 |
| NREM | 1.4 | 0.9 | 0.3 | 0.14 |
| REM | 0.8 | 0.5 | 0.6 | 0.44 |

### Milk Yield and Composition

There was no effect of treatment on milk production (*P =* 0.44; Table 6). Overall, the sleep deprivation treatment reduced protein content compared to lying deprivation (*P* = 0.01; Table 6). Lying deprivation tended increase SCC compared to sleep deprivation. However, there was no effect of day or treatment × day interaction (*P* = 0.64 and *P* = 0.15, respectively; Table 6).

**Table 6.** LS Mean and standard error of milk yield and milk components for lying and sleep deprived cows during 24 h baseline, and treatment days as well as the 4 days following treatment (excluding treatment day for milk production and days 2 to 4 for milk components). *P*-values reflect days compared to baseline.

| Variable | Milk Yield (kg) | *P*-value | Protein (%) | *P*-value | Fat (%) | *P*-value | SCC (cell/mL) | *P*-value |
| --- | --- | --- | --- | --- | --- | --- | --- | --- |
| Lying Deprivation |  |  |  |  |  |  |  |  |
| Baseline | 34.9 ± 2.5 | Ref | 2.9 ± 0.1 | Ref | 2.9 ± 0.2 | Ref | 53,596 | Ref |
| Treatment | -- | -- | 3.0 ± 0.1 | 0.04 | 3.6 ± 0.2 | 0.001 | 64,800 | 0.2 |
| Day 1 | 31.8 ± 2.5 | 0.001 | 3.0 ± 0.1 | 0.004 | 3.6 ± 0.2 | 0.001 | 61,464 | 0.4 |
| Day 2 | 32.8 ± 2.5 | 0.02 | -- |  | -- |  | -- |  |
| Day 3 | 34.6 ± 2.5 | 0.75 | -- |  | -- |  | -- |  |
| Day 4 | 36.2 ± 2.5 | 0.17 | -- |  | -- |  | -- |  |
| Sleep Deprivation |  |  |  |  |  |  |  |  |
| Baseline | 35.8 ± 2.5 | Ref | 2.9 ± 0.1 | Ref | 3.1 ± 0.2 | Ref | 53,873 | Ref |
| Treatment | -- | -- | 2.9 ± 0.1 | 0.6 | 3.6 ± 0.2 | 0.01 | 48,012 | 0.5 |
| Day 1 | 35.3 ± 2.5 | 0.61 | 2.9 ± 0.1 | 0.8 | 3.3 ± 0.2 | 0.14 | 41,159 | 0.08 |
| Day 2 | 34.1 ± 2.5 | 0.07 | -- |  | -- |  | -- |  |
| Day 3 | 36.0 ± 2.5 | 0.82 | -- |  | -- |  | -- |  |
| Day 4 | 35.2 ± 2.5 | 0.48 | -- |  | -- |  | -- |  |

When milk yield was compared between baseline (d −2) and the 4 days after treatment, cows produced less milk on d 1 and 2 (after trt) compared to baseline when deprived of lying (Table 6). Milk production tended to be lower on d 3 compared to baseline when sleep was deprived but did not differ on any other day relative to the baseline period.

When milk components were compared between baseline (d −2), treatment day, and then day 1 (day after trt), fat content was found to be higher on treatment day (d 0) and d 1 during lying deprivation, and on treatment day during sleep deprivation (*P* ≤ 0.01; Table 6). Protein content was higher on d 1, and 2 compared to baseline during lying deprivation (*P ≤* 0.04; Table 6). Protein content did not differ on any days during sleep deprivation (*P ≥* 0.05). For SCC, there was a tendency for a period and treatment effect to occur (*P* = 0.08 and *P* = 0.09, respectively).

## DISCUSSION

This study was the first to evaluate the effects of sleep and lying deprivation on behavior and milk production, which has not been well investigated in dairy cows. Previously, research has focused on the effects of lying deprivation, but has failed to consider the cumulative effect of lying and sleep deprivation during this time. Assessing the effects of sleep and lying deprivation separately is inherent to understanding the difference between gross quantity of lying time, and what she is doing while she is lying. Within the current study, both deprivations altered lying time after treatment, suggesting either deprivation could alter cow behavior and welfare. Furthermore, although sleep deprivation had no effect on milk production, the combination of sleep and lying deprivation reduced milk yield. All lying behaviors were lower on the day of the lying deprivation treatment compared to baseline. This finding is what we had expected, as our methodology was designed to eliminate lying time on the lying deprivation day. Our results and methodology were similar to previous studies that implemented lying deprivation (Metz, 1985, Munksgaard et al., 1999). Although we recorded a small amount of lying time during lying deprivation, the researchers who were present during the entire treatment period did not observe any lying time during the lying deprivation phase. This small amount of lying time was, thus, likely due to the loggers mis-reading cows that were shifting their weight to alleviate pressure on their hooves as ‘lying down’, as this has been recorded in other studies using similar loggers (Kok et al., 2015). We did not specifically observe these weight shifting behaviors as part of the study, but these behaviors have been observed as a sign of frustration and discomfort in other lying deprivation studies (Ruckebusch, 1975, Metz, 1985, Cooper et al., 2007). We are confident that the recorded lying time during lying deprivation was likely not real lying time.

Our sleep deprivation treatment did not reduce lying time relative to the baseline period. This suggests that cows remained lying down despite being roused awake by researchers. It should be noted that lying time during baseline for both treatments was less than previous reports for mattress bedding (Manninen et al., 2002, Tucker and Weary, 2004, Ito et al., 2009). Tucker and Weary (2004) reported a mean lying time of 12.3 ± 0.53 h/d on a mattress surface with no bedding. This may be due to the transition from the cow’s typical sand bedded freestall pen to an individual, mattress bedded pen, as cows change lying behaviors depending on bedding type (Tucker et al., 2003). Nonetheless, lying time the day after sleep deprivation was increased relative to treatment, suggesting some amount of lying time may be lost during sleep deprivation as well.

Lying bouts (4.9 ± 0.82 bouts/d) and bout duration (58.9 ± 7.31 min/bout) for both treatments during the baseline period were similar to previous literature, suggesting that researcher presence and the EEG equipment did not substantially disrupt lying behaviors. Previously, a mean of 8.5 ± 0.6 bouts/d (Tucker and Weary, 2004), and 10.7 ± 0.7 bouts/d (Manninen et al., 2002), were reported for dairy cows on mattress bedding. Although bout durations are shorter relative to reports by Tucker and Weary (2004) (90 ± 6.0 min/bout), data within the current study is similar to Van Gastelen et al. (2011), and Manninen et al. (2002), who reported 71.7 ± 10.2, and 70.4 ± 4.5 min/bout, respectively.

Lying bouts and bout duration had a tendency to differ between baseline and sleep deprivation, where during treatment, cows had a tendency to have less lying bouts and longer bout duration. However, lying bouts only differed by 1.4 bouts/d, and bout duration only differed by 11.09 min/d. Therefore, there may not be any biological relevance to the tendency, due to the minimal differences observed. Overall, lying deprivation altered lying bouts and bout duration more than sleep deprivation. Thus, cows can likely be sleep deprived without being fully lying deprived. However, the quality of that lying time during sleep deprivation is like reduced due to the inability to engage in sleep. Cows were previously estimated to engage in approximately 20 min of NREM sleep per day, 10 min of REM sleep, and 28 min of drowsing. Terman et al. (2012) reported bout lengths of 3 ± 2 min, 5 ± 3 min, and 3 ± 1 min for drowsing, NREM sleep, and REM sleep, respectively. Cows in the current study also engaged in relatively short duration bouts of these three vigilant states. Despite this consistency, there is the potential that the use of non-invasive techniques may lead to the underestimation of the vigilance states due to the physical characteristics, substantial skull thickness and the relatively modest brain size. The duration of rumination throughout the day as presents a challenge, as noted by Ternman et al (2018). Despite these potential issues, the results generated from non-invasive techniques do not differ substantially from Ruckebusch (1972), who utilized sensors implanted directly into the brain to assess vigilance state.

The number of steps taken within the current study differed depending on the study days. During the baseline period, cows took more steps than what was reported previously. For example, on sand bedded freestalls, cows took a mean of 1,611 ± 120.7 steps/d depending on the season (Kull et al., 2017). Although this is lower than steps taken within the current study, cow’s activity varies across environment and bedding type (Manninen et al., 2002, Tucker et al., 2003). Thus, cows were more active on the mattress bedding, relative to their normal sand bedded freestalls. Furthermore, number of steps taken during baseline for sleep and lying deprivation did not differ. This suggests that even though steps were higher than previously reported, baselines for both treatments were similar, indicating accurate comparisons can be made between treatments. The number of steps taken was less during baseline relative to lying deprivation, but did not differ between baseline and sleep deprivation. This implies that lying deprivation has a greater overall impact of a cow’s daily activity than sleep deprivation.

The EEG data provided further insight into the effects of treatment on the behavior of these cows. Previous work has varied considerably in the duration that cows were reported to be in various vigilant states. Early work estimated that cows spent 3 h/d in NREM sleep, 30 to 45 min/d in REM sleep, and 8 h/d drowsing (Ruckebusch, 1972); However, this is considerably longer than the 20 min/d in NREM sleep, 10 min per d of REM sleep, and 28 min per d of drowsing (Ternman et al., 2014) or 64 min per d of NREM, 44 min per d of REM, and 57 min per d of drowsing (Ternman et al., 2018) in more recent work on sleep in dairy cows. Cows in the current study were more closely associated with durations of NREM and REM sleep reported by Ternman et al. (2014) and Ternman et al. (2018), but closer to Ruckebusch (1972) for the duration of drowsing. The various vigilance states occur throughout the day in relatively short-duration bouts. Our results are consistent with Ternman et al. (2012) in this matter. As discussed previously, all studies of cows may underestimate true sleep time, especially for NREM, unless cows for some reason have a very atypical lack of neuronal synchronization leading to at least some NREM with high EEG delta power. However, this should not impact the current study, as sleep deprivation is never 100% complete. Drowsing may also have evolved as a light form of sleep and merits further investigation to understand the role this vigilance state has in the overall health and productivity of lactating cows.

The changes in vigilance states from baseline to treatment suggest that we were only partially successful in implementing our treatments. While we used the gentle handling technique to prevent cows from “sleeping” without depriving them of the ability to lie down, we did not effectively change the duration of time they spent in these various vigilance states during the treatment phase. This might have been driven by the poor agreement between observable behaviors and sleep in dairy cows (Ternman et al., 2014), which may have allowed these cows to engage in sleep without observable indicators. Conversely, our lying deprivation was effective at also inducing sleep deprivation. This begins to suggest that management activities, such as overstocking (Krawczel et al., 2012) or extended length of time restrained in headlocks (Cooper et al., 2007) that reduce lying times are also shifting sleep patterns. Further evaluations are needed to establish the changes in sleep caused by management factors and the role of this in productivity and feed efficiency.

Lying deprivation reduced milk production by 3.1 and 2.1 kg from the baseline period to d 1 and 2 (which are the 1^st^ two d after trt), respectively. Other studies either did not measure milk yield during lying deprivation (Ruckebusch, 1974, Metz, 1985, Munksgaard and Simonsen, 1996), or milk production was not affected (Munksgaard and Løvendahl, 1993, Cooper et al., 2007). However, when cows were lying deprived for 14 h/d for 23 d, growth hormone (GH) was reduced (Munksgaard and Løvendahl, 1993). GH in dairy cows is involved with the partitioning of energy resources in favor of milk production, as increased GH concentration is positively correlated with milk yield (Hart et al., 1978). While GH was not measured in the current study, it can be speculated that GH was a contributing factor to the reduction in milk yield for the lying deprived cows. Furthermore, GH hormone is also strongly tied to the sleep-wake cycle (Kim et al., 2011). GH secretion is typically increased during sleep, and suppressed during sleep deprivation (Brandenberger et al., 2000, Everson and Crowley, 2004). While, sleep deprivation did not have an effect on milk yield in the current study, it may be the cumulative effect of lying and sleep deprivation that reduced milk yield during lying deprivation.

Milk composition was altered during the experimental period. However, all components fell within the normal range. Within the current study, milk fat and protein were similar to other studies that reported a range from 2.0 to 6.1, and 2.5 to 2.8%, respectively (Kelsey et al., 2003). Furthermore, results are consistent with Åkerlind et al. (1999), and Bouraoui et al. (2002), who reported milk fat and protein similar to the results presented in the current study. Although milk fat was elevated during the treatment period relative to baseline, milk fat is the most variable of all components (Woodford et al., 1986), and changes based on lactation (Council, 1988), milking duration (Wheelock, 1980), and season (Jenness, 1985). Thus, milk fat changing slightly across days is not alarming, and may not be biologically relevant. Although feed intake was not measured in the current study, cows during lying deprivation do increase their feed intake (Cooper et al., 2007), which can increase milk fat percentage in dairy cows (Macmillan et al., 2017). This may be why fat content is higher during treatment and d 1, relative to baseline. SCC was increased for the lying deprived cows, relative to the sleep deprived cow. This may suggest that cows were stressed during this time, as SCC increased in cows during transportation (Yagi et al., 2004), and when mixed in groups (Kay et al., 1977), which can both be deemed as stressful events. However, SCC within this study were well below the 200,000 cell/mL threshold (Schepers et al., 1997, Bradley and Green, 2005), indicating the increase in SCC may not be biologically relevant. Collectively, the deprivation period may not have been long enough to alter milk composition significantly.

Overall, results were consistent among all lying behaviors for both treatments. The lack of differences between the baseline and treatment period for sleep deprivation suggests, while cows were sleep deprived, they were not lying deprived, indicating the successful separation of sleep and lying deprivation. While sleep deprivation alone did not reduce milk yield, there was likely a cumulative effect of lying and sleep deprivation during lying deprivation, and this may be why milk yield was reduced during lying deprivation. Furthermore, lying deprivation had a greater overall impact on cow activity and production.

### Treatment to Recovery Period Comparison

Lying time increased for both deprivations after the treatment period. However, on d 7, the last day of recovery, lying time was similar to previous research who observed a lying time of 9.5 to 12.9 h/d in freestalls (Ito et al., 2009), and 12.0 h/d on sand bedded freestalls (Cook et al., 2004). This suggests that d 7 may be more reflective of a cow’s typical lying time on sand bedding, and will be used as the comparison from this point forward. Lying time was higher on d 1 after lying deprivation, suggesting, lying deprivation strongly raises the need for lying (Metz, 1985, Munksgaard et al., 1999). The lying deprived cows lied down for longer on d 1, relative to the sleep deprived cows, indicating their need for lying may be stronger. Furthermore, it took the lying deprived cows 4 d to fully recover their lying time, whereas it only took the sleep deprived cows 1 d. This is likely due to the lying deprived cows losing more lying time during treatment, than the sleep-deprived cows. This speculation is further supported by cows who were lying deprived for 4 h/d, and lied down for longer during the post deprivation period, than cows who were deprived of 2 h/d (Cooper et al., 2007). This suggests, long lying deprivation periods result in higher lying times the subsequent days post deprivation. Overall, both sleep and lying deprivation increased lying time after deprivation, and therefore, altered a cow’s time budget and behavior. Thus, if her lying time is reduced due to lying or sleep deprivation, it could lead to poor welfare.

Lying bouts within the current study did not differ for either treatment, suggesting, cows did not have to recover any lying bouts after treatment. Additionally, results within the current study were consistent with prior data, who reported a range of 8.8 to 11.0 (Kull et al., 2017), and 10.2 to 10.3 bouts/d on sand bedding (Gomez and Cook, 2010). This indicated, even though lying time and bout duration were affected by treatment, the number of times cows lied down did not differ. To compensate for the loss in lying time, cows increased their bout duration rather than altering how many times they got up and down throughout the day.

Bout duration for the lying deprived cows was increased on d 1, relative to d 7, suggesting, cows lied down for longer before getting up the day after deprivation. However, bout duration during the rest of the recovery period was consistent with other studies who observed a mean of 88 (Ito et al., 2009), and 77 min/bout in a freestall environment. This increase in bout duration is likely driven by an increase in the motivation to lie from the lack of lying during treatment. However, since bout duration recovered after 24 h for the lying deprived cows, it is more easily recovered than overall lying time. Bout duration only had a tendency to differ on d 1 for the sleep deprived cows, suggesting, their lying time or bout duration was not as affect by sleep deprivation. Thus, bout duration for the lying deprived cows was altered more, relative to the sleep deprived cows.

Consistent with bout duration, steps followed a very similar pattern. Steps were consistent with the data presented by Kull et al. (2017), indicating, cows within the current study behaved similarly to other cows on sand bedding. However, steps did not differ for either treatment, the entire recovery period. This suggests, even though lying behaviors were altered, cows were likely taking the same number of steps/d, but spent less time standing idle, and more time lying, post deprivation. Typically, cows spend between 2.1 (Gomez and Cook, 2010) and 2.4 h/d standing idle (Cook, 2008), thus, this time was likely consumed by lying rather than standing.

While both deprivations altered behavior, lying deprivation may be the cumulative effect of both lying and sleep deprivation. This theory was first proposed by Moberg (2000), who believed the effects of multiple stressors being applied simultaneously, is biologically worse than experiencing one stressor. For example, when cows are heat stressed, their lying time is reduced as well (Cook et al., 2007, Herbut and Angrecka, 2017). While it is recognized that other physiological processes are altered during heat stress, the combination of both heat stress and lying deprivation, could be why other productive functions are also affected (Ravagnolo et al., 2000, Dash et al., 2016). Furthermore, when rats were restrained for a period of time, then injected with endotoxin, there were worse effects biologically, than the rats who only experienced only one of these stressors (Laugero and Moberg, 2000). This may be due to energy resources being shifted towards the stressor(s), and away from other productive functions such as growth (Moberg, 2000). Thus, the effects that occur during lying deprivation could be the cumulative effect of both lying, and sleep deprivation. This may be why worse effects on behavior are observed during lying deprivation, relative sleep deprivation.

In conclusion, both deprivations altered behavior after treatment. Thus, depriving cows of either sleep or lying long term may have worse effects than what was observed within the current study. Overall, lying deprivation had a greater impact on a cow’s lying time than sleep deprivation. Therefore, it may still be better for a cow to have access to an uncomfortable stall, where she can lie, but not necessarily engage in sleep, rather than not having a stall available at all. However, there is potential for cows to be experiencing both lying and sleep deprivation when lying time is reduced. Therefore, it could be the cumulative effect of both stressors occurring simultaneously, and why there are stronger changes in behavior during lying deprivation.

## CONCLUSIONS

Collectively, these data suggest that lying deprivation induces concurrent sleep deprivation, which has an additive, detrimental effect on the behavior and productivity, both yield and quality, of lactating dairy cows. This suggests that sleep deprivation may be one of the mechanisms explaining the decreases in productivity associated with reduced lying times. The limited effect on sleep resulting from the sleep deprivation treatment reinforces the hypothesis that cows can enter various vigilance states in a wide range of lying postures. This remains a critical question to address in order to evaluate management practices that reduce the potential quality of lying, such as poorly designed or managed lying areas, from those that reduce access to lying resources, such as overstocking or excessive use of headlocks or time in the holding area of milking parlors.

## ACKNOWLEDGEMENTS

This work was funded by a USDA-NIFA-Exploratory grant (USDA NIFA Exploratory Grant # 2015-67030-24295). The authors are grateful to the staff of the Little River Animal and Environmental Unit for the assistance with this project. The help of a large team of undergraduate students was also critical for the successful completion of data collection.

